# Transcriptome-wide association studies accounting for colocalization using Egger regression

**DOI:** 10.1101/223263

**Authors:** Richard Barfield, Helian Feng, Alexander Gusev, Lang Wu, Wei Zheng, Bogdan Pasaniuc, Peter Kraft

## Abstract

Integrating genome-wide association (GWAS) and expression quantitative trait locus (eQTL) data into transcriptome-wide association studies (TWAS) based on predicted expression can boost power to detect novel disease loci or pinpoint the susceptibility gene at a known disease locus. However, it is often the case that multiple eQTL genes colocalize at disease loci, making the identification of the true susceptibility gene challenging, due to confounding through linkage disequilibrium (LD). To distinguish between true susceptibility genes (where the genetic effect on phenotype is mediated through expression) and colocalization due to LD, we examine an extension of the Mendelian Randomization Egger regression method that allows for LD while only requiring summary association data for both GWAS and eQTL. We derive the standard TWAS approach in the context of Mendelian Randomization and show in simulations that the standard TWAS does not control Type I error for causal gene identification when eQTLs have pleiotropic or LD-confounded effects on disease. In contrast, LD Aware MR-Egger regression can control Type I error in this case while attaining similar power as other methods in situations where these provide valid tests. However, when the direct effects of genetic variants on traits are correlated with the eQTL associations, all of the methods we examined including LD Aware MR-Egger regression can have inflated Type I error. We illustrate these methods by integrating gene expression within a recent large-scale breast cancer GWAS to provide guidance on susceptibility gene identification.

## Introduction

Integrating data from genome-wide association studies (GWAS) of disease and expression quantitative trait loci (eQTL) studies can help detect novel disease loci and pinpoint genes of interest. Traditional studies have either matched the eQTL and GWAS association signals using ad-hoc overlap statistics or estimated the probability that the association signals are due to the same genetic variants (i.e., colocalization) [Giambartolomei, et al. 2014; Hormozdiari, et al. 2016; Wallace 2013]. More recent methods have proposed to test the association between transcript expression levels and disease risk by first using eQTL reference data to build multimarker predictors of expression, and then testing the association between the genetically predicted expression and disease risk in a large GWAS [Gamazon, et al. 2015; Gusev, et al. 2016a]. The latter methods, which we jointly refer to as Transcriptome-Wide Association Studies (TWAS), have been extended to the case when only summary statistics are available from the disease GWAS, the eQTL study, or both [Barbeira, et al. 2016; Gusev, et al. 2016a; Zhu, et al. 2016].

Although TWAS has been successful in identifying many genes whose genetically regulated expression is associated to traits, a major limitation of TWAS is that it cannot distinguish between a causal effect of expression on disease and a tagging association within the same region due to correlations among SNPs (i.e. linkage disequilibrium, LD) [Mancuso, et al. 2017a; Wainberg, et al. 2017]. A SNP used in expression prediction of gene A may be in linkage disequilibrium (LD) with nearby SNPs used in the prediction of gene B that is not involved in disease etiology. This LD will induce association between the genetically regulated expression of gene B and disease, causing the TWAS test statistic to reject the null of no association between predicted expression and disease at gene B even though expression levels are not causally related to disease; this effect is similar to standard LD-tagging in GWAS where LD induces significant association statistics at non-causal SNPs [Mancuso, et al. 2017a]. As TWAS methods were originally proposed as tests for association between local genetically regulated component of expression and disease with no causality guarantees [Gamazon, et al. 2015; Gusev, et al. 2016a; Mancuso, et al. 2017a; Mancuso, et al. 2017b; Zhu, et al. 2016], it remains unclear whether and when TWAS can be interpreted as valid tests of causality. In contrast to TWAS, colocalization analyses, including COLOC, eCAVIAR, focus on estimating the probability of the SNPs causals for eQTL to be the same as for GWAS, irrespective of direction of genetic effect on expression or disease, and are not designed to test for an effect of gene expression on disease risk [Giambartolomei, et al. 2014; Hormozdiari, et al. 2016; Wallace 2013].

In this work, we explore the use of TWAS methods in the context of causal gene identification. As opposed to the standard TWAS that mainly focused on association testing to identify new genomic regions harboring disease genes, in this work we investigate the utility of TWAS methods in the context of causal gene localization (which utilizes a much more stringent definition of true/false positive). We re-derive TWAS methods in a Mendelian Randomization (MR) framework and show that the standard TWAS statistic is a special case of MR that use eQTLs as genetic instruments to test the causal association between expression and disease. We leverage the growing literature on methods for MR that use summary statistics for both the associations between single nucleotide polymorphisms (SNPs) and the intermediate trait and between SNPs and disease; that use multi-SNP genetic instruments [Burgess, et al. 2016]; and that remain valid when some of the assumptions underlying standard MR are violated [Bowden, et al. 2015]. In particular, the MR Egger regression approach relaxes the assumption that the association between genetic instruments and disease is only mediated through the intermediate trait—which would not be the case if the SNPs in the genetic instrument had pleiotropic effects, for example [Bowden, et al. 2015]. We investigate the use of a variant of MR Egger regression that accounts for LD among the variants used in the genetic predictor for gene expression (LD aware MR Egger regression) in the context of TWAS. This method was first proposed independently of this group in the discussion section of Burgess and Thompson 2017 with derivations provided in their appendix [Burgess and Thompson 2017]. We use extensive simulations to compare the performance of LD aware MR Egger to other TWAS and MR statistics and show that it remains valid under specific violations of the assumptions underlying MR, while retaining comparable power to the other approaches when the assumptions do hold.

The structure of this paper is as follows. First, we introduce the conceptual model relating the outcome and gene expression to the SNPs. Second, we discuss four existing approaches for testing the association between the outcome and gene expression. We show that the traditional TWAS test statistic of Gusev et al. [Gusev, et al. 2016a] is equivalent to an LD aware version of standard MR using summary statistics and compare this approach to LD aware MR-Egger regression (LDA MR-Egger) [Burgess and Thompson 2017]. Third, we examine the statistical properties of these estimates and their performance via simulation in the presence or absence of a direct effect of SNPs on disease. To the best of our knowledge, this is the first time the empirical performance of LDA MR-Egger regression has been examined in this context. We apply the various approaches to summary statistics from a GWAS on Breast Cancer [Michailidou, et al. 2017] with eQTL data from a breast tissue panel in GTEx [Consortium 2013]. We conclude by providing guidance on the interpretability of standard TWAS and MR tests can be interpreted as valid tests in the context of eQTL and GWAS integration.

## Methods

Main Models

Let ***Y*** denote the outcome (*nx1*), ***M*** the mediator (*nx1*), and ***G*** the SNP matrix (*n*x*J*) of interest. We assume that the columns of ***G*** have been standardized to have mean zero and variance one. In the TWAS setting, ***M*** is gene expression. We denote the LD structure of the SNPs ***G*** as ***Σ***, a *JxJ* symmetric positive definite matrix. (In practice, this matrix may be a modified version of the empirical LD matrix, adding a small ridge regularization to ensure the matrix is symmetric positive definite and invertible [Pasaniuc and Price 2017]). Here we let ***Σ*** represent the correlation with Σ_*k,j*_ represent the Pearson correlation between SNP k and SNP j. To motivate our approach, we assume that (for a link function *g*):

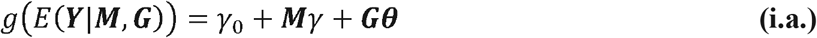

and

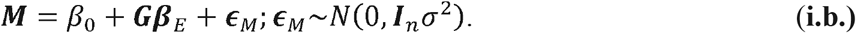

We are interested in the solution to the regression of ***Y*** on ***G***, marginal over ***M***:

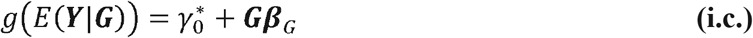

In the above models, ***θ*** is a *J*-column vector of ***G*** effects on ***Y*** conditional on ***M**, γ* is the effect of ***M*** on ***Y*** conditional on ***G, β**_E_* is the *J*-column vector of SNP effects on the mediator, and ***β**_G_* is the *J*-vector of the ***G*** effects marginal over ***M ϵ**_M_* and ***I**_n_* represent the residual variance in ***M*** and the *n* x *n* identity matrix respectively. We are interested in the situation where we cannot directly estimate the parameters in model (*i.a*), as we do not have complete data on ***Y, M*** and ***G*** from (sufficiently many) individuals. We want to place inference on *γ*, because if *γ* ≠ 0, gene expression ***M*** affects the trait. To relate the parameters in the marginal model (1.c) to the models (1.a.) and (1.b.), we assume one of the following for the remainder of the paper:

1. *g* is either the log or identity link function.
2. **Y** is a sufficiently rare binary trait and *g* is the logit link.

If either of the two conditions above hold, we will have:

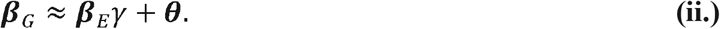

If *g* is the log or linear link, the approximation will be exact. Equation (ii) suggests that the effects of ***G*** marginal over ***M*** are a function of the eQTL parameters (***β**_E_*), the effect of the gene expression on the outcome conditional on ***G*** (*γ*), and the effect of ***G*** on the outcome conditional on *M*(***θ***). We will call ***θ*** the direct effect and ***β**_E_γ* the mediated effect, but we note that ***θ*** will be non-zero not only when ***G*** has has a direct causal effect on ***Y***(i.e. not mediated through ***M***), but also when the relationship between ***G*** and ***Y*** is confounded—as would be the case when there is colocalization and the SNP(s) with an effect on the outcome are not included in ***G***. We will explore what happens in the later scenario empirically.

In practice, we do not have an overlap of data on ***M, Y***, and ***G***, and hence cannot estimate *γ* directly. Instead, we have a sample of size *N* that the GWAS is run on to estimate ***β**_G_* and an independent sample of size *N_E_* that was used to estimate ***β**_E_*. Moreover, we typically only have estimates of individual SNP effects marginal over the other SNPs, 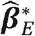 and 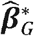. For our purposes, 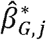 and 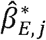 were estimated with the same reference allele for SNP *j*. If they were not, the sign of the effect can be changed so as to refer to the same reference allele. We note here that while in this paper we refer to ***β**_G_* as the GWAS effects, these parameters can be estimated from any association study.

The formulas given above relating the mean of ***Y*** and ***M*** to ***G*** were given on the conditional level. We therefore transform our marginal estimates (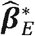 and 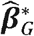) to the conditional scale. Given an estimate of the LD matrix (***Σ***) we estimate the conditional eQTL and GWAS effects as 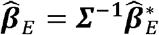 and 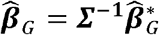 [Shi, et al. 2016]. If the marginal effect estimates were not calculated on the standardized genotypes they can be transformed by multiplying by the square root of 2*p_j_*(1-*p_j_*), where *p_j_* is the MAF of SNP *j*. We assume that 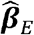 and 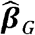 are unbiased for ***β**_E_* and ***β**_G_* respectively.

We do not know the form of ***θ***; it could be a constant or vary by SNP. All we know is that it is a vector of length *J*. Our estimated GWAS effects, (given our assumptions above) are a function of ***θ, β**_E_, γ* and the sampling error:

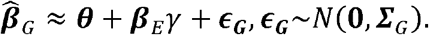

If the SNPs are not in LD, then the marginal and the conditional will be equal 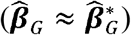.

We next derive the covariance of 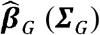. Let 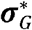 denote a *Jx1* vector of the marginal standard errors of 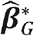. Let “.” denote element wise multiplication between two matrices. As ***G*** has been standardized, 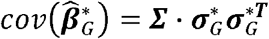. This gives the covariance of our conditional GWAS estimates:

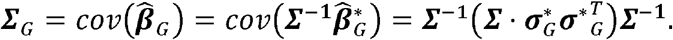

If 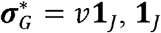 is a column vector of ones, then 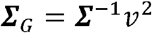. The main goal is to derive a valid test of the null hypothesis *γ*=0 and potentially estimate *γ*in the presence of direct effects of the SNPs onto the outcome. We next go over some common methods for testing for an association between gene expression and outcome.

### Transcriptome Wide Association Studies (TWAS)

The standard TWAS statistic uses summary statistics to test for an association between genetically predicted gene expression and a phenotype of interest [Gusev, et al. 2016a]. The TWAS does not necessarily estimate the *γ* above but does provide a valid test for the association between the gene of interest and the outcome. The TWAS test statistic is:

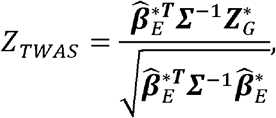

where 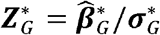, is a column vector of the marginal test statistics. The test statistic is then compared to a standard normal to assess significance. The TWAS can use either the marginal or conditional eQTL estimates as weights. If using the conditional, the equation above becomes:

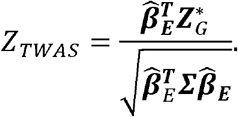

### The Summary Mendelian Randomization Estimator

The summary MR estimator [Johnson 2011] is:

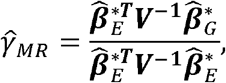

where ***V*** is a diagonal matrix with 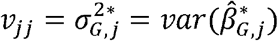. If there are no direct effects, i.e. ***θ*** = 0, and the SNPs are not in LD with each other (***Σ*** = ***I**_J_*), then this will be an unbiased estimate of *γ*. The MR estimate is best suited for analysis where the mediator is another phenotype and the SNPs are taken from across the genome. This estimate (which we term as the MR estimate) can be rewritten as:

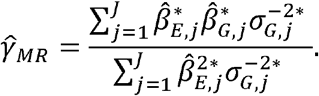

We estimate its variance as:

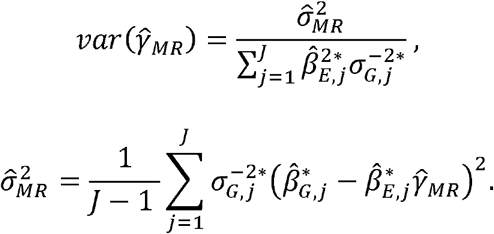

The MR test statistic is then:

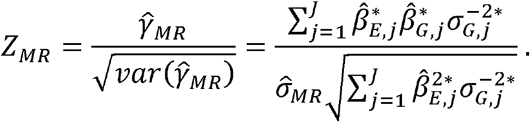

For testing, we compare *Z*_MR_ to the quantiles of a *t*-distribution with *J*-1 degrees of freedom. If the SNPs are in LD or if there are direct effects, 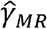 can lead to incorrect inference. The MR estimate is a weighted linear regression without an intercept of the marginal GWAS estimates on the marginal eQTL estimates with weights equal to 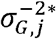.

We note here that Zhu et al. refer to their single SNP MR test as a summary Mendelian Randomization, that approach is not the same as what is presented here [Zhu, et al. 2016]. The approach above takes into account all SNPs at once as opposed to doing one SNP at a time. The Zhu et al approach provides an estimate for pleiotropy (HEIDI) based on the summary statistics and conditioning on the lead SNP.

### MR-Egger Estimate

If the MR estimate is a weighted linear regression without an intercept, the MR-Egger extends the MR by including an intercept (*α*) to the weighted linear regression. It assumes that 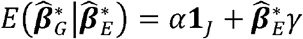. The estimates are (same ***V*** as the MR estimate):

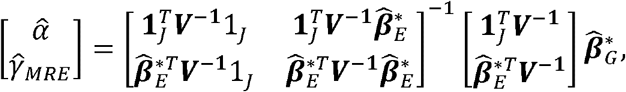

The estimate of *γ*is then:

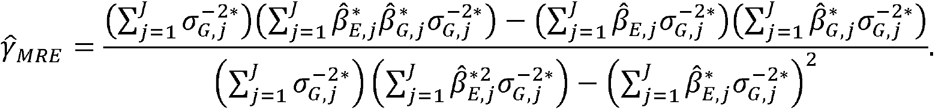

The variance of the estimate is:

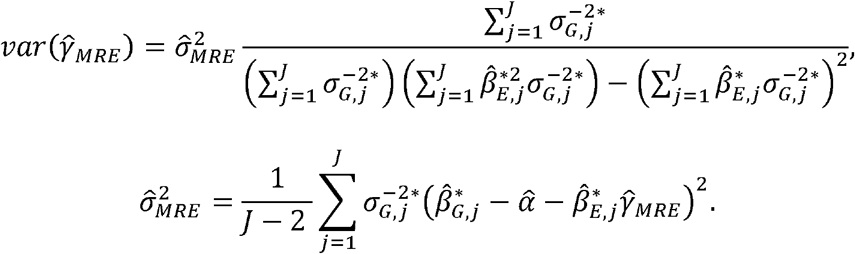

The MR-Egger test statistic is then:

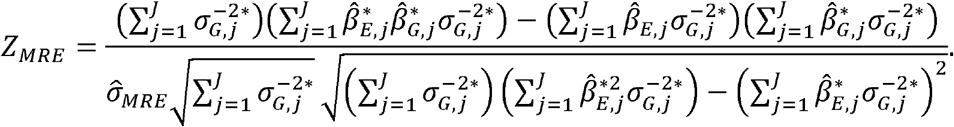

To test, we compare to the quantiles of a *t*-distribution with *J*-2 degrees of freedom. If the SNPs are in LD, the test for 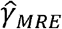 may lead to incorrect inference due to the variance of 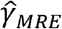 being misspecified.

### LD-Aware MR (LDA MR) Estimate

The LDA MR estimator of *γ* extends the MR estimator by relaxing the assumption of the SNPs being independent. It however, still requires that there are no direct effects. Recall that 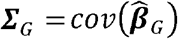. The LDA MR estimator is then:

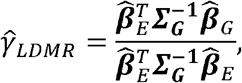

And we estimate the variance as:

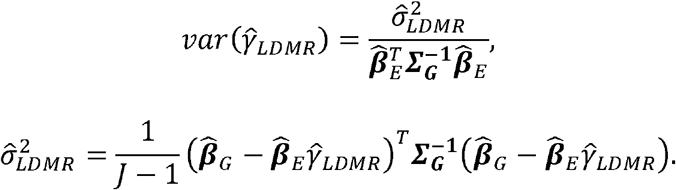

The test statistic is then:

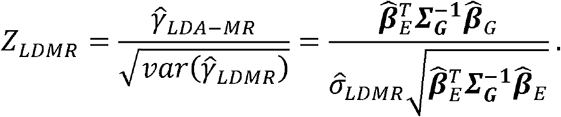

Similar to the MR, we compare to a *t* distribution with *J*-1 df. If there are direct effects, 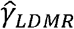 can lead to incorrect inference. Just as the MR estimate is a weighted linear regression without an intercept, the LDA MR estimate is a weighted linear regression without an intercept using the weight matrix 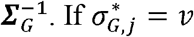 for all SNPs, the LDA MR and the TWAS test-statistic run on the marginal eQTL will be proportional to each other by 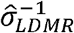 (Appendix A).

### LDA MR-Egger Estimate

LDA MR-Egger regression extends the MR-Egger approach to incorporate the LD structure of the SNPs. We include an intercept, in the aims of accounting for the direct effect of the SNPs. The estimates of the intercept and *γ* are:

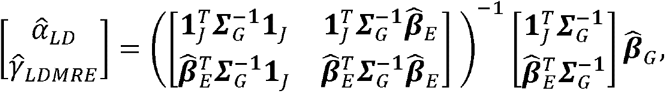

with *γ* more succinctly being written as:

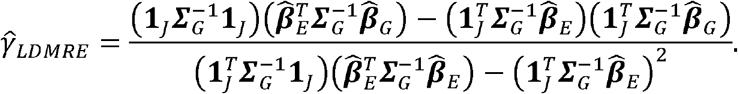

The variance is estimated as:

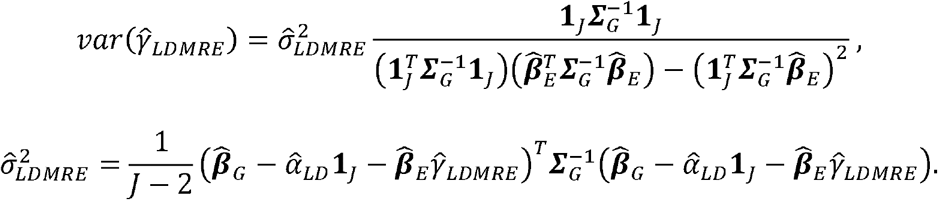

The test statistic is then:

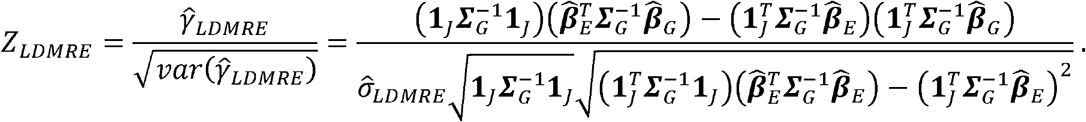

For testing, we compare to a *t*-distribution with *J*-2 degrees of freedom. This estimate is a combination of the approaches from LDA MR and the MR-Egger. The LDA MR-Egger will provide valid inference in the same scenarios as the MR-Egger, but in addition when the SNPs are in LD. Details of all tests are provided in Table I.

### Biases of Estimates

We first focus on the estimates that do not incorporate an intercept, the MR and the LDA MR. We now examine the bias of the MR estimate:

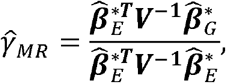

Note that 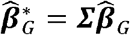 and 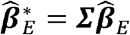. Thus 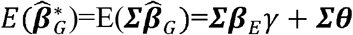. We assume that 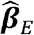 and 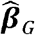 were estimated from different samples and thus are independent, 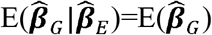. Using this gives us that:

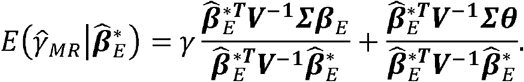

Then use that 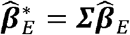:

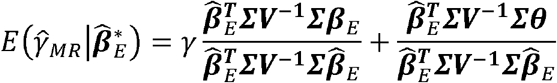

The term 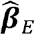 was estimated from a sample size of N_E_. As N_E_ →∞, we again have that 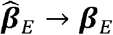 and the first term goes to *γ* Assuming N_E_ is sufficiently large:

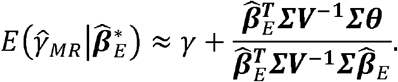

Now unless ***θ*** = 0 or the transformation 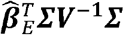 is orthogonal to ***θ***, the MR will be biased. For the LDA MR:

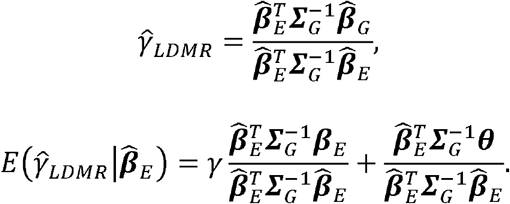

As 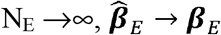 and the first term will go to *γ*. Assuming that N_E_ is sufficiently large enough for this to occur, we have:

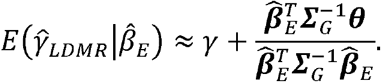

Again, if ***θ*** ≠ 0 or if the transformation 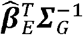 is not orthogonal to ***θ***, then the estimate will be biased. As *J* increases, the N_E_ needed for 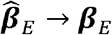 will also increase.

We next assessed methods that include an intercept. For the MR-Egger estimate, using the fact that 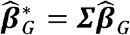 and 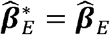, we have:

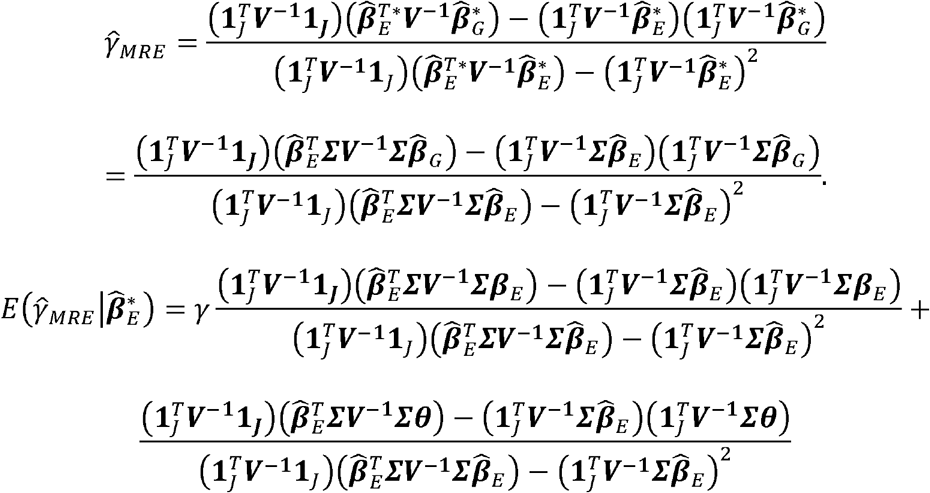

As we have done previously, we assume N_E_ is sufficiently large so that 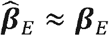, making the first term *γ*.

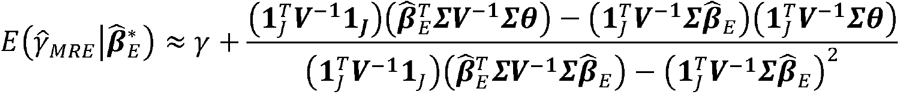

Thus, the MR-Egger estimate will be unbiased if there is no direct effect (***θ*** = 0), or if ***θ*** is a constant, or if the numerator in the second term is equal to 0. Note that if ***Γ***=***I*** and 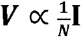, as is the case in the setting where the MR-Egger estimate was originally proposed, then the numerator is equal to the empirical covariance between 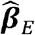 and ***θ***. This condition is referred to as the Instrument Strength Independent of Direct Effect (InSIDE) condition [Bowden, et al. 2015]. If the number of SNPs in the instrument for the mediator (*J*) is large, and there is no systematic relationship between ***β**_E_* and ***θ***, then this empirical covariance will be small. However, if *J* is small, as may be the case when gene expression is the mediator and the genetic instrument is limited to cis SNPs, then the numerator in the second term may be non-zero even when there is no systematic relationship between ***β**_E_* and ***θ***. Even if the INSIDE condition holds but ***Σ*** ≠ **I**, the MR-Egger estimate will lead to improper inference, as it does not account for linkage disequilibrium.

We now consider the LDA MR-Egger estimate:

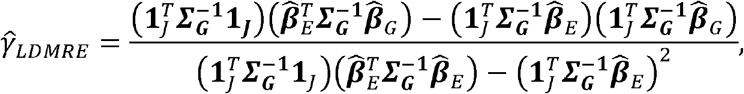

Again, using that 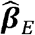 and 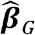 were estimated from different studies, we have that

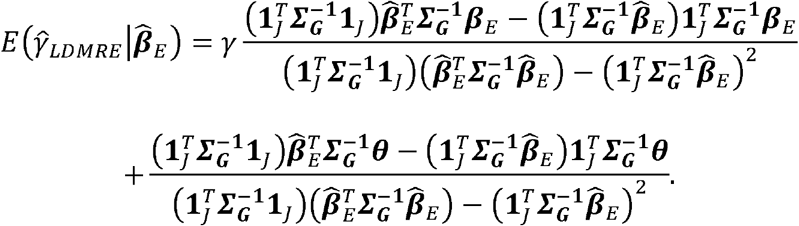

As with the other estimates: 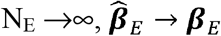. If we assume that N_E_ is sufficiently large:

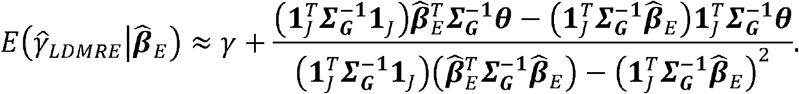

The numerator of the second term is a function of the sample univariate covariance between 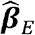 and ***θ*** weighted by 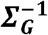. The LDA MR-Egger estimate will be unbiased in the same situations as the MR-Egger estimate, but it does not require the SNPs be independent.

### Simulation Study

We examine the empirical performance of four methods: the MR, MR-Egger, LDA MR, and the LDA MR-Egger. We note here that the MR and LDA MR approaches’ empirical performance in the presence of LD has been examined in detail previously [Burgess, et al. 2016], but for the sake of comparison we include them in our analysis. Under our simulation scenario, the TWAS and the LDA MR test statistics are approximately the same (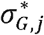 is a constant), so we only present results on LDA MR. We generated the data such that all SNPs have a small causal effect. For each simulation, we generated summary eQTL and outcome SNP statistics from a multivariate normal distribution as opposed to individual level data [Han, et al. 2009]. We fixed the sample of the eQTL study to 1000 (N_E_) and the sample size of the GWAS to 5000 (N). We varied the number of SNPs at the locus (*J*); the proportion of variation in ***Y*** explained by 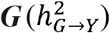 and 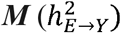; the proportion of variation in ***M*** explained by 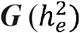; the magnitude of any shared direct effect (*τ*); and the LD matrix (***Σ***, AR(*J*) structure, *Σ_i,j_* = *ρ*^|*i*−*j*|^). For all simulations, the expression effects ***β**_E_* and direct effects ***θ*** were sampled independently. More details along with values taken are given in Table 2. For a variable direct effect, *τ* was set to 0. For a near-constant, shared direct effect, *τ* = 10. When *τ* = 10 the direct effects of SNPs on disease are all in the same direction (*θ_i_* > 0 for every SNP *i*) and the variability in *θ* in this case is about 1% of the variability when *τ* = 0. *τ* can be thought of as a shrinkage parameter, with a larger *τ* decreasing the variability in the direct effects (***θ***) and shrinking towards a common effect. We refer to this situation as “directional pleiotropy.” The process and order for the generation of the simulation data is provided in Table 3. We performed 50K simulation for each combination of parameters (486 different combinations). In each simulation, we generated a new true ***β**_E_* and ***θ***, which are functions of 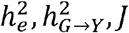 and *τ*. Generating this new “true” parameter value better represents the different eQTL and GWAS patterns across different genes, and therefore more resembles a standard TWAS. The procedure detailed in Table 3 is thus repeated 50K times for all combinations of the parameters in Table 2. Type I error (T1E) and power were evaluated at 0.05.

**Table 1:** Commonly used approaches for testing using summary statistics. Details four common Summary Mendelian Randomization approaches and the TWAS for testing for an association between gene expression and outcome through GWAS (*J* is the number of SNPs in the loci).

| Method | Adjusts for Direct Effect | Accounts for LD | Distribution |
| --- | --- | --- | --- |
| TWAS | No | Yes | $Z_{TWAS} \sim N(0,1)$ |
| MR | No | No | $Z_{MR} \sim t(df = J - 1)$ |
| MR-Egger | Yes | No | $Z_{MRE} \sim t(df = J - 2)$ |
| LDA MR | No | Yes | $Z_{LDMR} \sim t(df = J - 1)$ |
| LDA MR-Egger | Yes | Yes | $Z_{LDMRE} \sim t(df = J - 2)$ |

**Table 2:** Simulation parameters that were modified. Parameters that were varied for all simulations performed. A description of the parameter and potential values could be taken are given. All combinations of parameters were examined.

| Parameter | Definition | Values taken |
| --- | --- | --- |
| $h_e^2$ | Proportion of variability in E explained by G | 0.01, 0.20, 0.50 |
| $h_{E \rightarrow Y}^2$ | Proportion of Variability in Y explained by E | 0, 0.005, 0.01 |
| $h_{G \rightarrow Y}^2$ | Proportion of Variability in Y explained by G | 0, 0.005, 0.01 |
| $N$ | Sample Size of GWAS | 5000 |
| $N_E$ | Sample Size of EQTL | 1000 |
| $\rho$ | Correlation between SNPs. AR structure | 0.125, 0.9 |
| $J$ | Number of SNPs in the Loci | 50,300 |
| $\tau$ | Strength of Pleiotropic Effects | 0,10 |
| $\mathbf{L}$ | Cholesky Decomposition of $\Sigma$ | Function of $\rho$ |

**Table 3:** Procedure for generating simulation study. Gives the steps in order for generating the type I error and power simulations detailed in the paper and how the parameters were generated.

| Step | Procedure | Mathematically |
| --- | --- | --- |
| 1 | Generate true eQTL | $\beta_E \sim N_J(0, \mathbf{I}_J)$ |
| 2 | Set the proportion of variability<br>in expression due to SNPs | $\sigma_E = \sqrt{\frac{h_e^2}{\beta_E^T \Sigma \beta_E}}$ |
| 3 | Rescale the eQTL effects | $\beta_E = \sigma_E \beta_E$ |
| 4 | Set Expression to outcome<br>Effect | $\gamma = \sqrt{h_{E \rightarrow Y}^2}$ |
| 5 | Generate potential Direct<br>Effect | $\theta = e + \tau; e \sim N_J(0, \mathbf{I}_J)$ |
| 6 | Set the proportion of variability<br>in outcome due directly to<br>SNPs | $\sigma = \sqrt{\frac{h_{G \rightarrow Y}^2}{\theta^T \Sigma \theta}}$ |
| 7 | Rescale the Direct Effects | $\theta = \sigma * \theta$ |
| 8 | Generate the GWAS effects | $\beta_G = \theta + \gamma \beta_E$ |
| 9 | Generate Observed eQTL Values | $\hat{\beta}_E^* \sim \Sigma \beta_E + L^T \epsilon_E; \epsilon_E \sim N_J \left( 0, \frac{1 - h_E^2}{N_E} I_J \right)$ |
| 10 | Generate Observed GWAS values | $\hat{\beta}_G^* \sim \Sigma \beta_G + L^T \epsilon_G; \epsilon_G \sim N_J \left( 0, I_J \frac{1 - h_E^2 * h_{E \rightarrow Y}^2 - h_{G \rightarrow Y}^2}{N} \right)$ |

We in addition performed two more sets of simulations to solely assess the T1E. The first scenario, we calculated the empirical T1E under a non-infinitesimal model for both disease and expression (i.e. when only a subset of SNPs were associated with either trait). Under this scenario, SNPs were randomly assigned to be either an eQTL SNP, a disease SNP, or neither. We generated under the scenario where 10% (or 50%) of the SNPs were eQTL SNPs, and 10% (or 50%) of the SNPs were disease SNPs. The LD between these two sets would naturally vary as the SNPs were randomly assigned to their classification. We evaluated the T1E under the same scenarios as above, performing 50K simulations and evaluating at 0.05. This scenario represents the situation when there is colocalization of eQTL and disease SNPs within the same locus, but all SNPs are included in the analysis. We next simulated when the InSIDE condition was violated. To do so, we varied the correlation between *θ_j_* and *θ_E,j_* to be 0.125, 0.5, or 0.9. This correlation between direct and expression effects could result when there are disease SNPs nearby that are in LD with the SNPs included in the model (colocalization).

As ***β**_E_* and ***θ*** were drawn independently, the bias for the LDA MR-Egger estimates across many independent simulations of ***β**_E_* and ***θ*** should be 0. (This will also be true for the MR-Egger estimates in the absence of LD.) This is analogous to saying the bias across all the genes in the genome will be zero. However, in practice, we will usually be interested in the test statistic applied to a particular gene. For particular gene with modest *J*, the bias for the LDA MR-Egger need not be 0. To account for this, we next performed a set of 10K simulations with a fixed truth to examine potential bias. We generate one true ***β**_E_* and ***θ*** for each value of *J* that is then held constant for all simulations while we vary the other parameters in Table 2. Therefore, Steps 1 and 5 of Table 3 are only performed once for J=50 or 300. We also compared our estimate of *γ* to when a regularization factor of 0.1 is added to the diagonal of the LD matrix as mentioned in the beginning of the methods [Pasaniuc and Price 2017].

### Application to Breast Cancer GWAS Summary Data

We next applied TWAS, LDA MR, and LDA-MR Egger analyses applied to a breast cancer GWAS. We previously conducted a separate breast cancer TWAS using a different approach to build expression weights and including both validation of predicted expression and functional follow up of significant genes [Wu, et al. 2017]. Here our focus is to compare different analysis approaches where each analysis uses the same set of simple expression weights and the same GWAS summary statistics. These expression weights are different from that of the Wu et al paper [Wu, et al. 2017]. The marginal GWAS summary statistics were from a recent GWAS on breast cancer within women of European descent [Michailidou, et al. 2017]. SNP data was meta-analyzed across 13 GWAS; more details can be found in Michailidou et al. [Michailidou, et al. 2017]. After QC, the study consisted of 11.8 million SNPs, with 105,974 controls and 122,977 cases. The GWAS estimates were calculated on the non-standardized minor allele counts, and therefore were transformed using the minor allele frequency.

Expression weights were calculated from GTEx along with LD information in breast tissue in an overall sample of 183 individuals [Consortium 2013]. We restricted our analysis to the set of transcripts that were deemed heritable using GCTA [Yang, et al. 2011] and examined SNPs within 500kb of the gene boundary. The expression weights were calculated on standardize minor allele counts of SNPs (mean zero, variance one) and were conditionally estimated using the BSLMM approach [Zhou, et al. 2013]. A gene was deemed heritable if the GCTA p-value for each tissue from GTEx was less than the Bonferroni threshold of 0.05 (after adjusting for 27,945 tests). We were left with 683 transcripts to analyze.

We analyze the Breast Cancer data for these genes using the TWAS, LDA MR, and LDA MR-Egger. We did not examine the MR and MR-Egger as we were testing for cis-signals as opposed to genome wide and therefore the SNPs were in LD. We took the overlap of the 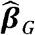 SNPs from the GWAS with the available SNP correlations from GTEx. If the effects were estimated with respect to a different reference allele, we reversed the sign for that eQTL effect estimate. In total, the 683 genes corresponded to 191,583 unique SNPs.

## Results

### Simulation Study

First, we examined the type I error rate in simulations and observed that when there is little LD and no direct effect of the SNPs, all of the approaches have the correct type I error (Figure 1, Sup Figure 1). When there is little LD and the direct effect is variable, the MR and MR-Egger approaches have modestly inflated Type I Error rates. When there is low LD and there is directional pleiotropy, only the LDA MR-Egger has correct type I error. If the SNPs are in high LD (bottom row of Figure 1) and there is no direct effect, the MR and the MR-Egger have inflated type I error due to misspecification of the variance. When there is a variable direct effect, all four approaches have inflated type I error. Finally, when there is strong linkage and directional pleiotropy, only the LDA MR-Egger has the correct type I error.

**Figure 1:**
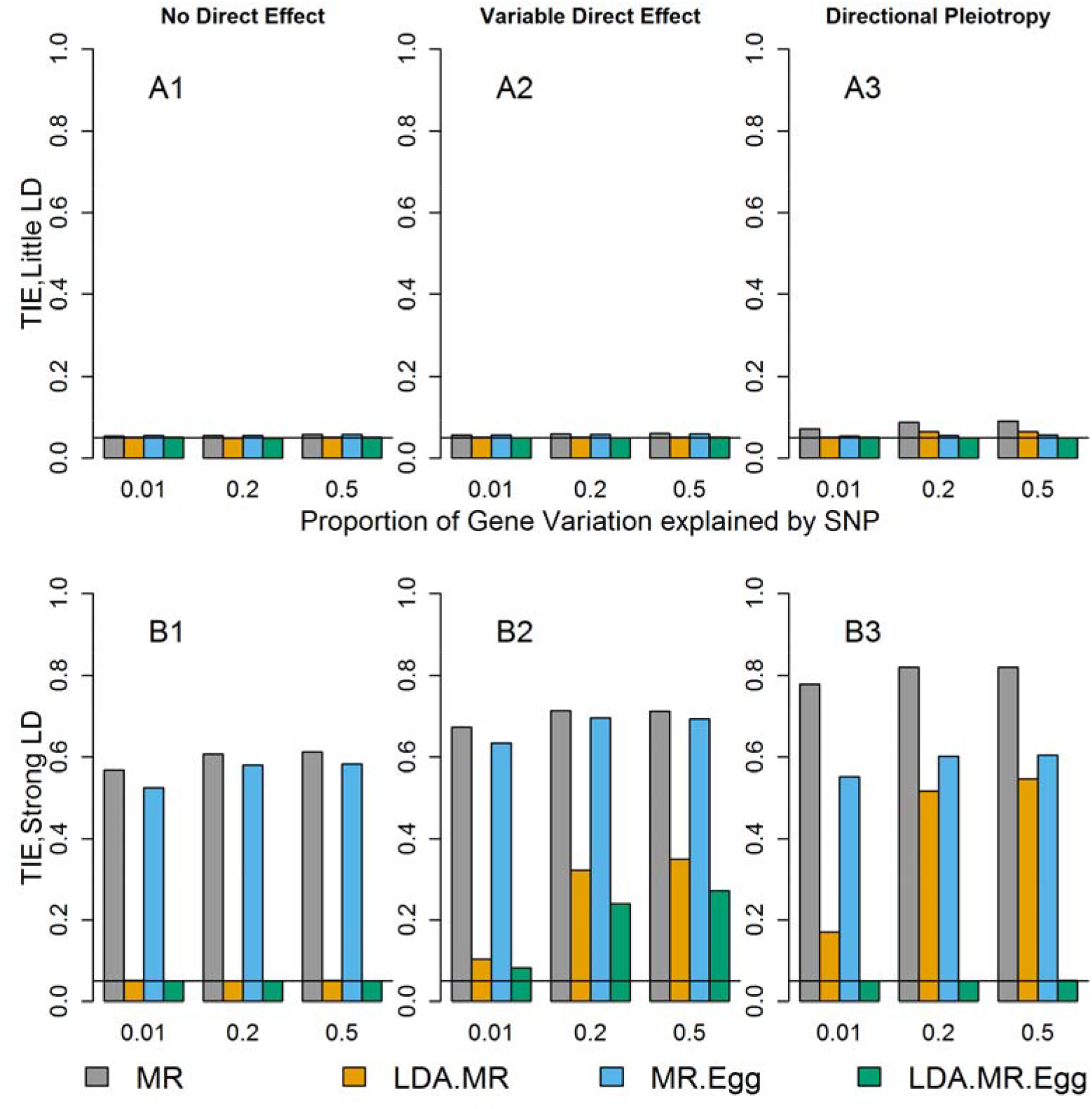
Type I error when J=50. Each bar represents results over 5×10^4^ simulations. Evaluated at *α* = 0.05. First panel represent when low LD (plots with A). Second panel represents when strong LD (plots with B). From left to right correspond to: no direct effect, variable direct effects with mean 0 across SNPs, and direct effects with mean >0 across SNPs and small variability (directional pleiotropy). When there is a direct effect, the SNPs explain 1% of the variation in the outcome 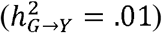.

We next examined the power when there is little to no LD between the SNPs and no direct effect (Figure 2). Under this situation, all four approaches had correct type I error (Figure 1 top left plot, Sup Figure 1 top left plot) and are valid tests. Regardless of the magnitude of the effect of **M** on **Y**, we see similar power for all four methods. There is slightly smaller power for the LDA MR-Egger compared to the LDA MR or MR when the SNPs only explain 20% of the variation in the gene expression, but once the SNPs explain 50% of the variation in ***M***, all approaches have approximately equal power. In each individual plot, we see that as the SNPs explain more of the variation (and are thus better instruments), we have an increase in power. It has been previously reported that the MR Egger regression can have notably lower power than MR [Bowden, et al. 2015]. We did not observe large differences in power, but we consider situations with more SNPs in the genetic instrument for the intermediate factor and large GWAS sample sizes. (For example, simulations in Bowden et al. (2015) consider 25 SNPs up to 1,000 subjects in the GWAS.) We performed additional simulations assuming only 5 SNPs were included in the genetic instrument; in this case, we also observed lower power for the LDA MR Egger method relative to the LDA MR method.

**Figure 2:**
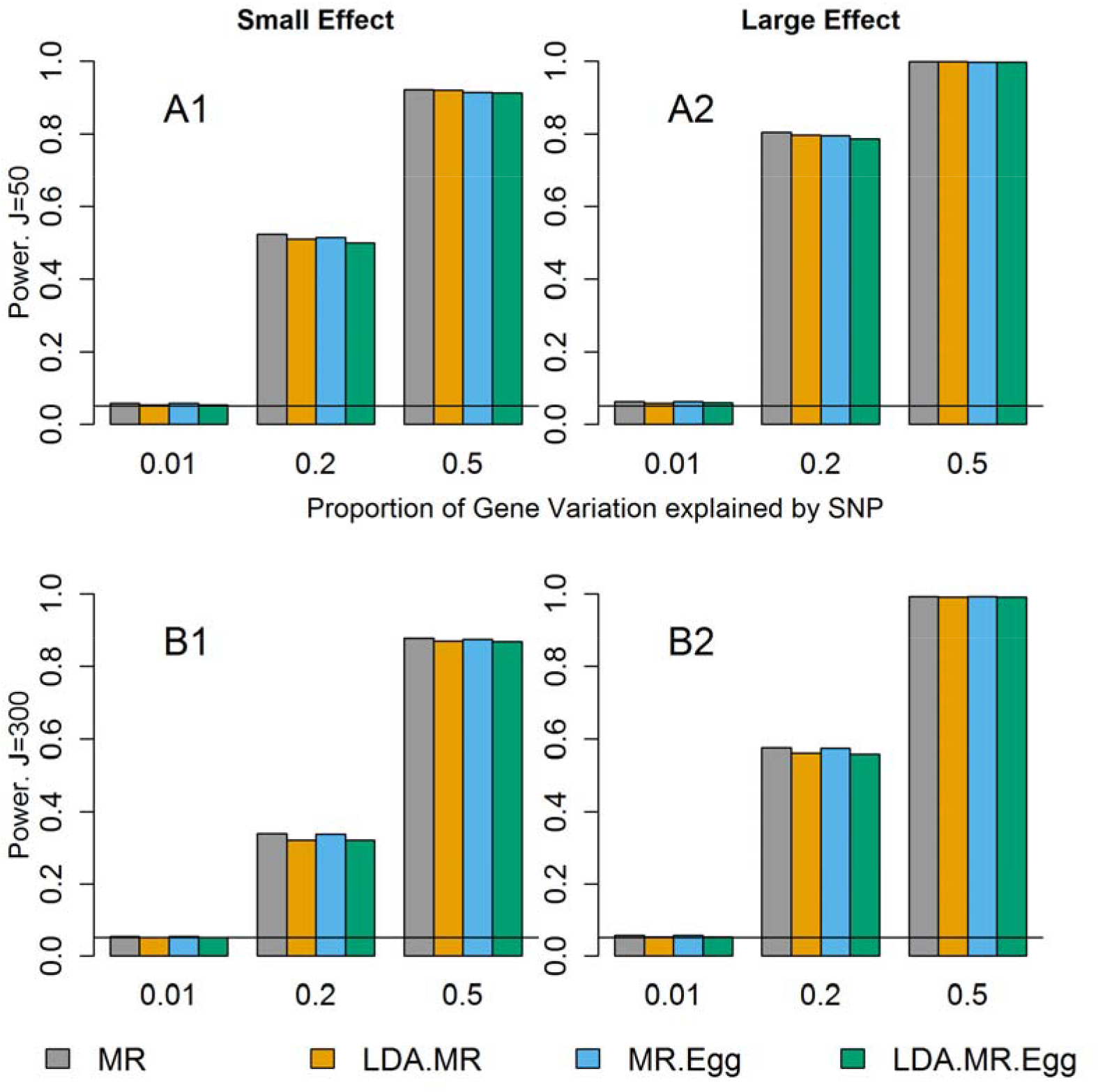
Power when little to no LD and J=50, 300. Power results when there is little to no LD and no direct effect. Each bar represents results over 5×10^4^ simulations. Evaluated at *α* = 0.05. First row represents J=50 and second row when J=300. From left to right: when γ^2^ = 0.005 and γ^2^ = 0.01.

The decrease in power from J=50 to J=300 is due to an increase in the noise to signal ratio when predicting expression levels using SNPs. For J=50, a larger proportion of the variation explained by SNPs is shared by each individual SNP leading to more precise estimates. When J=300, a smaller proportion of that same amount of variation is explained by each SNP in a larger set of SNPs. Assuming the sample size in the reference panel used to estimate 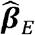 is the same, the sampling error in the SNP-specific estimates 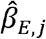 is the same for J=50 and J=300. We have held the proportion of variance explained by the SNPs constant, while increasing the noise due to sampling error (as we are using estimated expression effects from 300 SNPs rather than 50).

When there is low LD and directional pleiotropy (Table 4), then the MR-Egger approach has slightly higher power than the LDA MR-Egger test to detect an association when J=50. If J=300, there is a decrease in power compared to when J=50 for both of these methods regardless of the presence of a direct effect. At J=300, the LDA MR-Egger has slightly lower power than the MR-Egger. When the SNPs explain 50% of the variation in ***M***, both methods have power greater than 80% regardless of the effect of ***M*** on ***Y*** (Table 4). We did not report the power of the MR or LDA MR, as they did not have proper type I error when there is directional pleiotropy (Figure 1 and Supplement Figure 1).

**Table 4:**
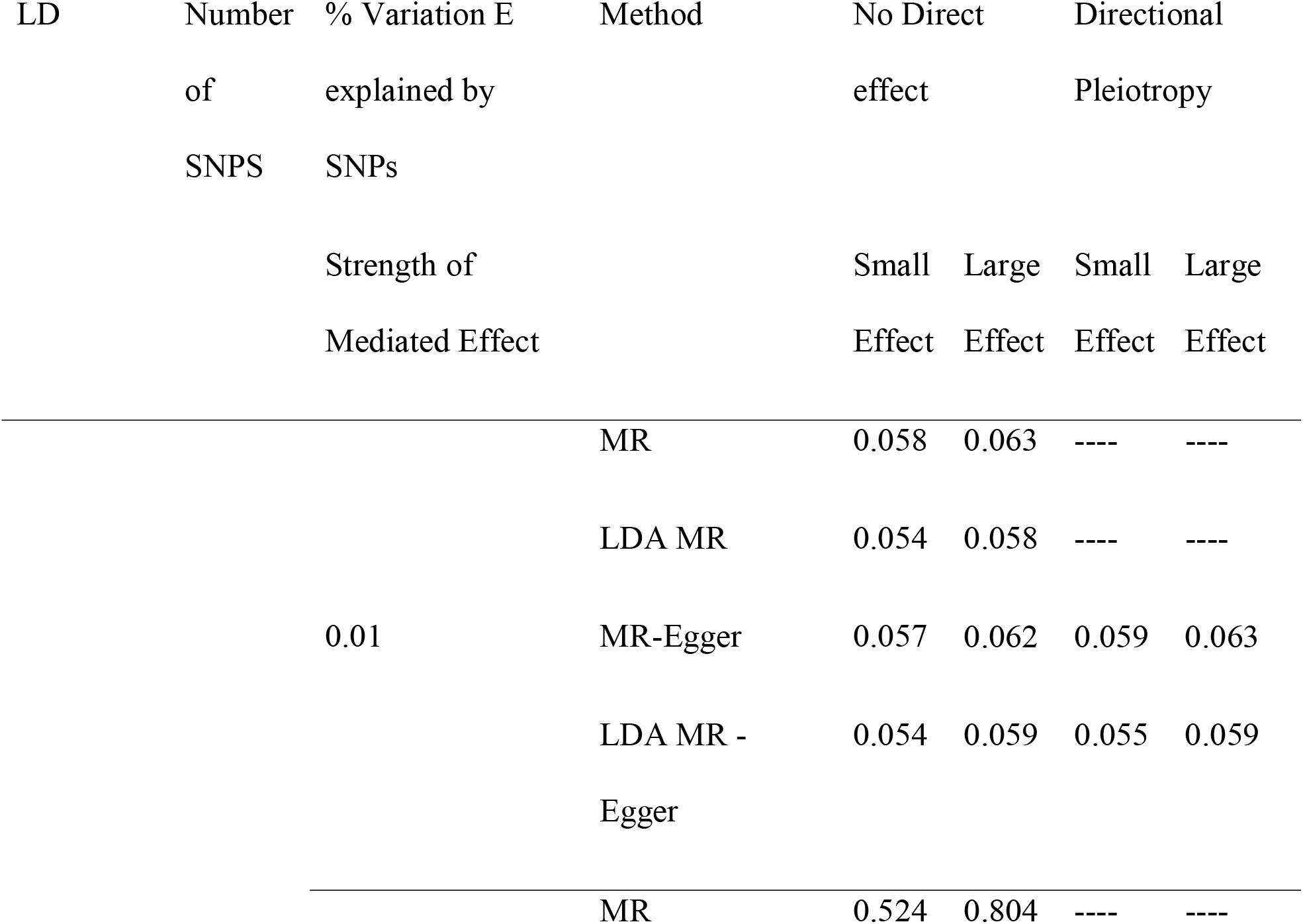
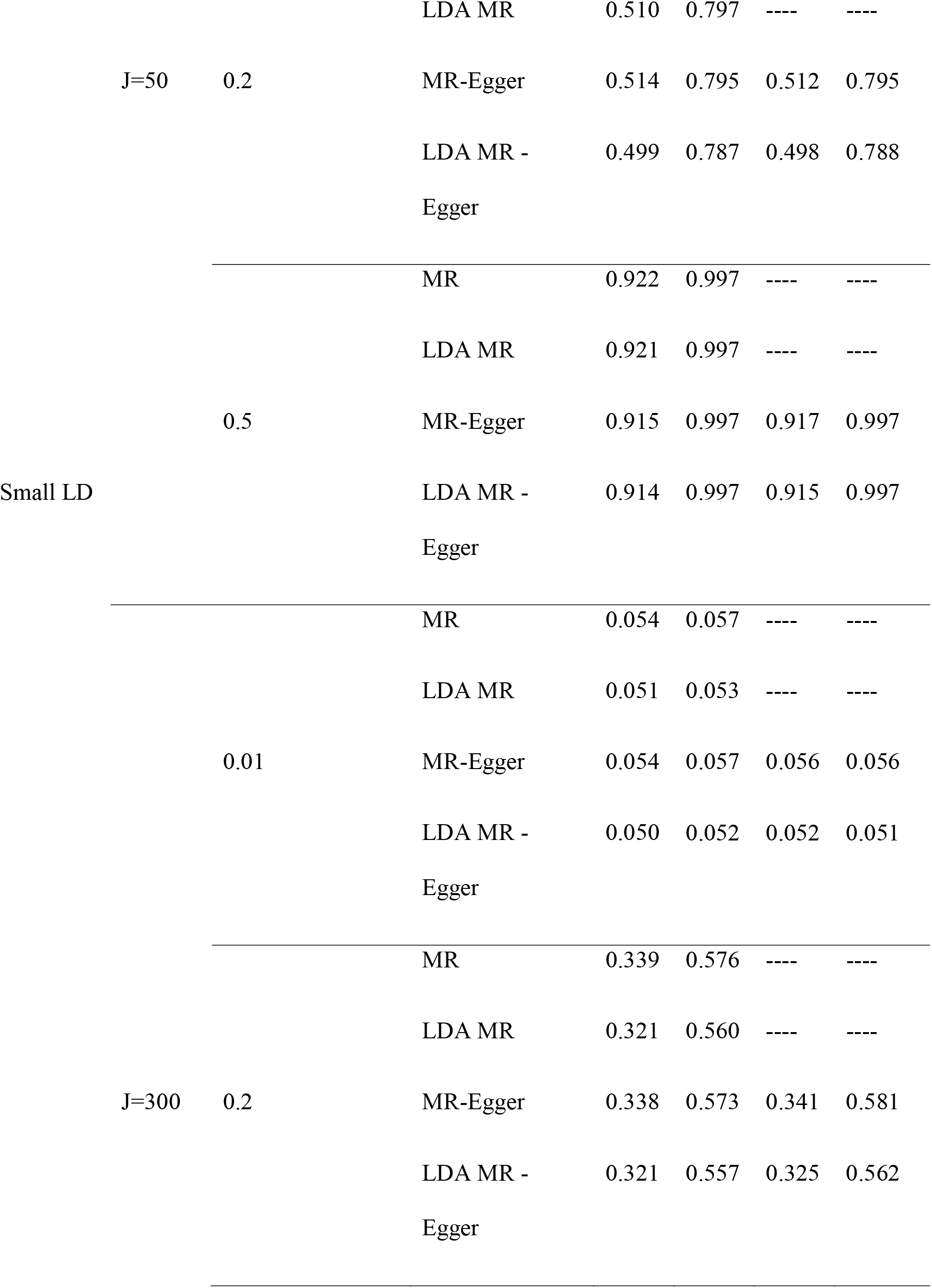
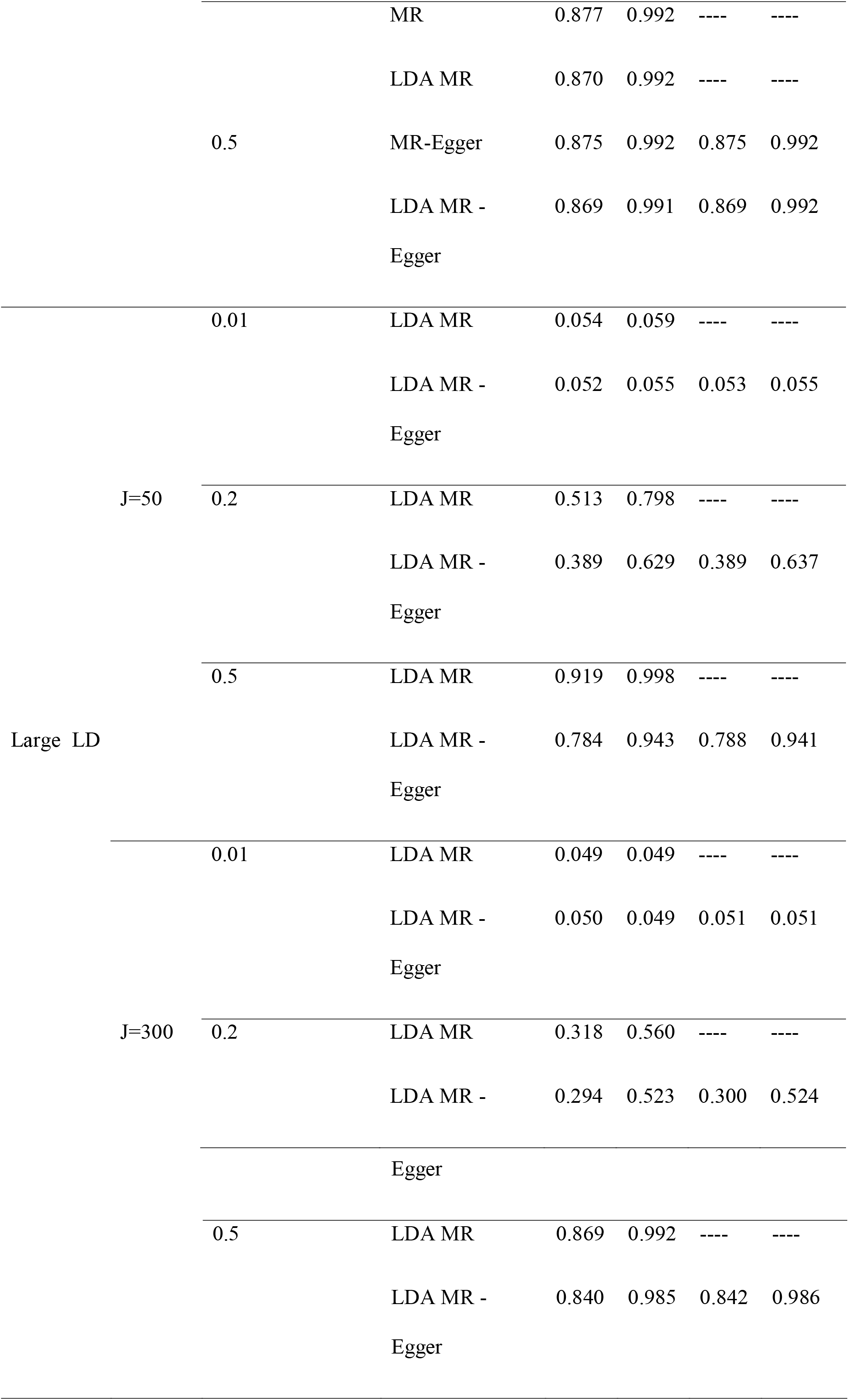
Power results from simulation study. Details the power results at an *α* level of 0.05 under varying levels of LD, number of SNPs in the loci, and presence of a direct effect. TWAS not shown as is equivalent to the LDA MR under are simulation procedure.

Finally, we examine the power when there is strong LD amongst the SNPs. We do not assess the MR or MR-Egger for this scenario as they do not have correct type I error for correlated SNPs. When J=50, and there is no direct effect, the LDA MR has more power than the LDA MR-Egger, though the difference in power is less pronounced when 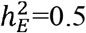 and ***M*** has a strong effect on ***Y***. When J=300, the two methods have comparable power (Table 4), with the LDA MR-Egger having slightly less power than the LDA MR. When there is a direct effect, the LDA MR-Egger has approximately equal power as when there is no direct effect.

When J=50, 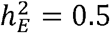, and there is a small mediated effect, then LDA MR-Egger regression has more power when there is strong LD versus low LD (power of 0.914 versus 0.784). When J=300, there is a smaller drop in power for small LD to strong LD, with the LDA MR-Egger power going from 0.869 to 0.840. The LDA MR approach has equal power regardless of whether the SNPs are strongly correlated vs weakly correlated.

The results for the non-infinitesimal simulations are provided in Supplementary Figures 2-5. For the most part, these results mirror what was observed in the standard T1E analysis. The only difference was when J=50 and there was little LD (Supplement Figure 2). When there is a variable direct effect and the SNPs explain an adequate amount of the variation in gene expression, we see all four approaches are conservative. In this case, where a small number of SNPs have relatively large effects on expression, the residual errors in the regression of GWAS effects on eQTL weights may be increased, which in turn will deflate the test statistics.

When the InSIDE conditions are violated, we see that regardless of if the direct effect are near-constant or variable around 0 we have massively inflated T1E (Figure 3 and variable direct effect Supplement Figure 6). The results for when J=300 are given in the supplement and are similar (Supplement Figures 7 and 8). This inflated T1E was expected given the violation of the InSIDE condition. This violation could happen when the true disease SNPs are in LD with SNPs in the model but are not included in the model (colocalization). This could also potentially happen if the SNPs influence a third trait that then goes on to affect the outcome and is correlated with the mediator.

**Figure 3:**
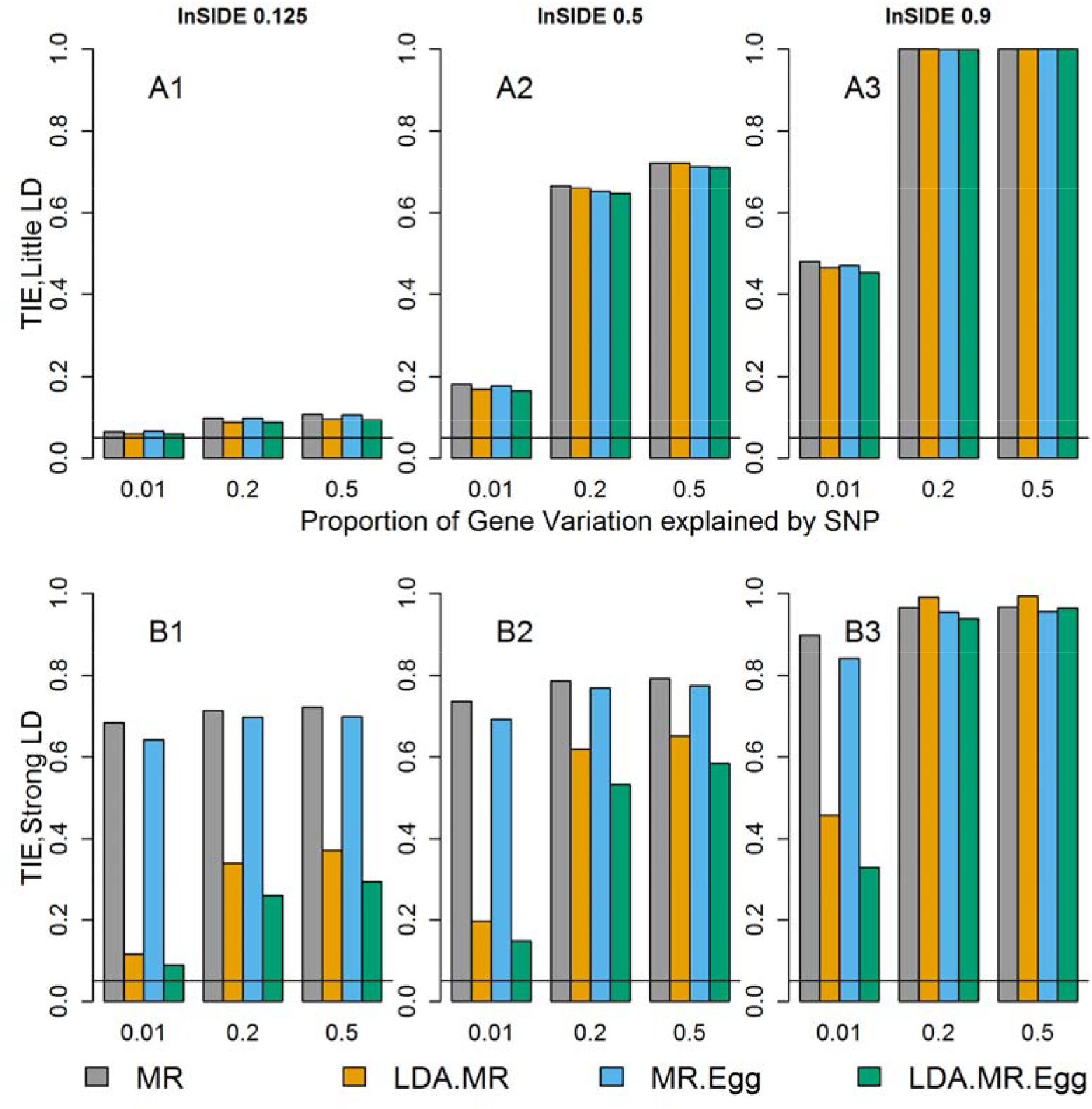
Type I error when J=50, InSIDE condition violated, and there is directional pleiotropy. Each bar represents results over 5×10^4^ simulations. Evaluated at *α* = 0.05. First panel represent when low LD (plots with A). Second panel represents when strong LD (plots with B). From left to right correspond to: correlation between *θ_j_* and *β_E,j_* is 0.125, 0.5, or 0.9.

We next examine the bias of our estimates (Figure 4). We here show the results when there is strong LD (Figure 4). The results for low are in LD Supplement Figure 9. In Figure 4, the first two rows show the results when J=50, and the next two rows when J=300. The first column shows when there is no direct effect, the second column when there is a variable direct effect, and the third column when there is directional pleiotropy. The first and third row are when there is no effect of the mediator on the outcome (*γ* = 0) and the second and fourth row show when there is a large effect (*γ* ≠ 0). Here we have fixed ***β**_E_* and ***θ*** effects for all simulations.

**Figure 4:**
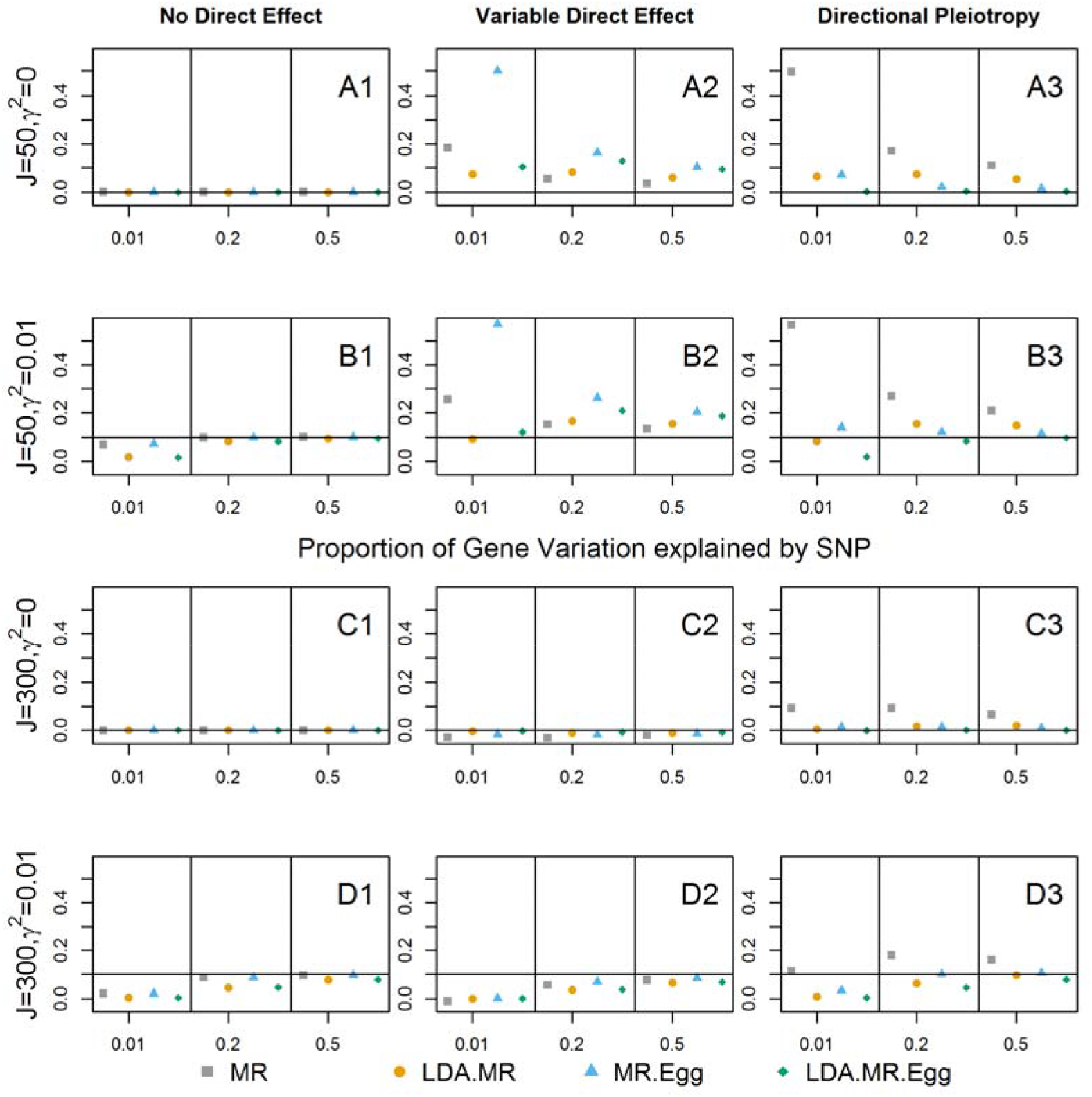
Bias when strong LD for J=50, 300. Bias plots for when there is strong LD in the SNP set. First row corresponds to J=50, γ = 0 (plots with A). Second panel (plots with B) when J = 50 and γ^2^ = 0.01. Third panel (plots with C) when J = 300 and *γ* = 0. Final panel (plots with D) J = 300 and γ^2^ = 0.01. From left to right: no direct effect, variable direct effects with mean 0 across SNPs, and directional pleiotropy. When there is a direct effect, the SNPs explain 1% of the variation in the outcome 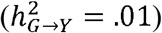.

As expected based on the section on bias of estimates sections, when *γ* = 0 and ***θ*** = **0**, all of the estimates are unbiased. When there is an effect on the outcome and no direct effect, we see that the non LDA aware approaches converge faster to the truth than the LD aware methods. While the non LDA converge faster, recall that they misspecify the variance and lead to improper inference. We also see attenuation bias when *γ* ≠ 0, with estimates improving as the SNPs become better instruments. The attenuation bias is larger for J=300 relative to J=50, for the same reason that we saw a decrease in power from J=50 to J=300: a decrease in the signal to noise ratio. When there is a variable direct effect, all of the approaches are biased (second column). Our empirical bias result depends on the particular values of ***β**_E_* and ***θ***. As described in the methods (“Bias”), all of the estimates for the mediated effect of gene expression include a weighted covariance between ***β**_E_* and ***θ***. Even if the average across all genes for this covariance is zero, for any particular gene it is non-zero, and can be large. If the ***β**_E_* and ***θ*** for each gene can be thought of as independent draws from their respective distributions, then the average absolute magnitude of the bias term decreases from J=50 to J=300. This is why we see smaller bias when *J*=300. Finally, when there is directional pleiotropy, only the LDA MR-Egger is unbiased when *γ* = 0 or J=50. When J=300 and *γ* ≠ 0, we see the attenuation bias in the LDA-MR Egger, with it biased downward toward the null. We saw little difference between the LDA methods based on adding a regularization constant of 0.1 to the diagonal or not of the LD matrix (Supplement Figure 10-11).

### Application to Breast Cancer GWAS Summary Data

Of the 683 genes tested, 79 were called significant (p< 7.32*10^−5^) by at least one approach (TWAS, LDA MR or LDA MR-Egger, Supplementary Tables). Comparing the TWAS vs the LDA MR-Egger (Figure 5A), there were 12 genes that were significant by the TWAS but not by LDA MR-Egger, 26 genes that were by LDA MR-Egger and not TWAS, and 20 that were called by both. With the LDA MR and the LDA MR-Egger, there was much more agreement due to the same weight matrix being used, but still the LDA MR called 8 genes as significant that the LDA-MR Egger did not (Figure 5B). Thirty-eight gene transcripts were called significant by both the LDA-MR and the LDA-MR Egger methods (Supplement Tables 1 and 3). There were 19 genes called significant by all three approaches. A detailed list of which gene transcripts were found significant by which method is provided in Supplementary Tables 2 through 8. Examining the spearman correlation between the p-values, LDA MR and LDA MR–Egger had an r^2^ of 0.45, LDA MR and TWAS of .51, and LDA MR-Egger and TWAS of 0.31. The kappa statistic for calling a gene transcript significant between LDA MR and LDA MR-Egger was 0.63, between LDA MR and TWAS was 0.55, and between LDA MR-Egger and TWAS was 0.48. We note that although the TWAS and LDA MR statistics are equivalent when the GWAS effect estimates have constant variance (see methods), in practice the GWAS effect estimates will differ across SNPs (e.g. due to sample size differences), leading to the differences in test statistics we see here. The Pearson correlation squared between TWAS and LDA MR test-statistics was 0.72 (Supplement Figure 12).

**Figure 5:**
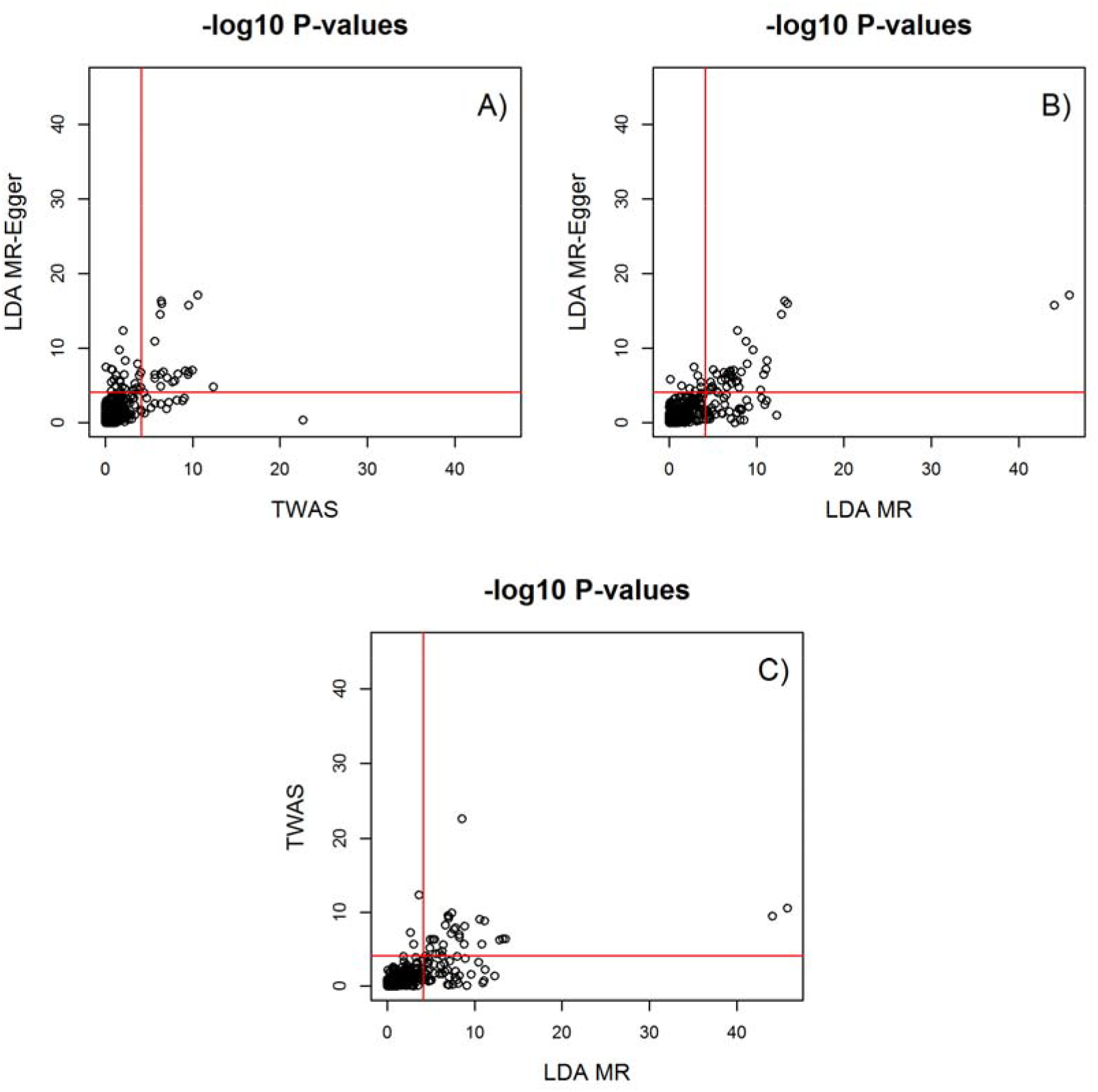
Comparing −log 10 p-values. Shows for 683 genes between LDA MR Egger and TWAS (A), LDA MR Egger and LDA MR (B), and LDA MR and TWAS (C). Red line is the Bonferroni cutoff of −log_10_(.05/683).

A gene that we will highlight that was called significant by the TWAS was SET Domain Containing 9 (*SETD9*), with a p-value of 2.47e-23 at cytoband 5q11.2. The p-value for this gene transcript for the LDA MR and the LDA MR-Egger was 2.97e-09 and 0.41 respectively. This is potentially a TWAS and LDA-MR false-positive as the LDA MR-Egger was not associated with gene. A fine mapping analysis of this locus found four functional candidate SNPs in a sample of approximately 100K women of European descent [Glubb, et al. 2015]. These four candidate functional SNPs were associated with an increase in activity of *MAP3K1* (Mitogen-Activated Protein Kinase 1), another gene at this locus. *SETD9* was ruled as not the gene of interest as it had no association with these four candidate SNPs. *MAP3K1* is located 94 kbp from *SETD9*. (*MAP3K1* did not pass our cis-heritable tissue threshold and was not included in the analysis.)

## Discussion

In this work we connect TWAS with Mendelian Randomization (MR) and examine the performance of the LD aware MR-Egger regression in the context of causal gene expression identification. In contrast to TWAS that mainly focused on novel risk region identification, here we investigate causal gene identification at a given risk region, a more difficult problem; for example, traditional TWAS does not aim to disentangle between two genes that show significant associations at the same risk region (e.g., due to LD-tagging between SNPs in the expression prediction models for the two genes). We show that the standard TWAS method is a particular case of MR. We investigate a variant of MR that allows for LD (LD aware MR-Egger) that can account for LD and direct effects under the scenario when the variability in direct effects across SNPs is small and the InSIDE condition holds. The LDA MR, the summary MR and the MR-Egger are not proper estimates under these scenarios as they either do not account for: the direct effect (LDA MR, MR) or the correlation between SNPs (MR and MR-Egger). For scenarios where the LDA MR is a valid test, the LDA MR-Egger has slightly lower power in our simulations. We also provide a one-to-one relationship between TWAS and the traditional LDA-MR when the standard errors of the marginal GWAS effects are all equal. In our real data application, the three approaches (TWAS, LDA MR and LDA MR-Egger) called different genes significant, with the LDA MR-Egger potentially correctly calling some results as null that the TWAS did not. While we focused on the case of gene expression data, the LDA MR-Egger can be extended to an arbitrary mediator when the instruments are correlated.

The majority of our simulations assumed an “infinitesimal” genetic architecture for both the gene expression and outcome phenotypes. Departures from this model—for example, if gene expression is causally influenced by only one or a small proportion of local SNPs not included in the model—can affect the performance of the tests. We saw this in our simulations where the InSIDE condition was violated. Future work could consider the impact of local genetic architecture on these tests. We mainly focused on the case where there may be pleiotropic direct effects on the outcome, but these are not systematically related to the SNPs’ effects on the mediator. This is reflected in our simulations when we draw ***β**_E_* and ***θ*** independently. If there is a systematic relationship between ***β**_E_* and ***θ***, then none of the methods we have discussed here will provide unbiased estimates of *γ* or valid tests of *γ* = 0 (Figure 3). There could be a systematic relationship between ***β**_E_* and ***θ*** even when *γ* = 0 if, for example, the SNPs influence a third (unobserved) trait, which in turn influences both the mediator and outcome, or if there is a confounder of the ***G-Y*** relationship. Of particular concern, the InSIDE condition could be violated as a result of colocaliztion, if the analyzed eQTL SNPs are in LD with an unmeasured (or unanalyzed) disease SNP. As many TWAS approaches select a subset of SNPs at a locus when building a genetic instrument for gene expression, it is possible that the SNPs included in a TWAS analysis do not include nearby disease SNPs. One method that might ameliorate this problem would be to use a model for gene expression prediction that does not involve SNP selection, such as ridge regression, and then apply LDA MR-Egger. Another would be to condition the TWAS test statistic on individual SNPs that are associated with outcome but not included in the expression prediction model[Gusev, et al. 2016b; Yang, et al. 2012]. How well these methods can control inflated T1E due to colocalization requires further investigation. Evaluating the causal effect of the mediator on outcome when the instrument and direct effects are systematically related may not be possible without additional information on the mediator-outcome relationship: large samples with data on genetic factors, the mediator, outcome and possible confounders will likely be needed.

In practice, we note that meaningful biological interpretation of the magnitude of *γ* may be difficult, due to QC and pre-processing (scaling) of the data. (The sign of *γ*, however, may contain useful information regarding the directionality of expression effects on disease.) We are also faced with the issue of attenuation bias, as we have estimates of the true eQTL parameters. Despite this, the LDA MR-Egger is still a valid test for the effect of the gene on outcome if the required assumptions are met. To reiterate these assumptions: they require that the variability in the direct effect of the SNPs on disease is relatively small and that the mechanism that the direct effect acts through is independent of the eQTL effect (InSIDE condition). The LDA MR-Egger can still however help to highlight a region of interest. Practitioners should follow up any analyses with a deep literature search paired with examining functional information such as pathways, enhancers, and other tissue expression. Whereas traditional epidemiological MR studies need to be certain of the causal pathways, the goal of these methods is to identify target genes whose expression levels are likely to influence disease risk. These genes represent candidates for functional experiments in model systems to investigate the effects of perturbing gene expression.

In summary, we have examined the performance of various summary statistics approaches and how they compare to the LDA MR-Egger. The LDA MR-Egger approach can be utilized only under the following situations: 1) No Direct Effect, 2) There is limited variability in direct effects of the SNPs on the outcome and this effect is independent of the eQTL effect. If we are not in one of those scenarios, the LDA MR-Egger test for the gene expression’s effect on outcome will not be valid—although neither will any of the other tests we considered This work provides guidance on the interpretation of TWAS tests and suggests that LDA MR-Egger regression may be a useful sensitivity analysis in situations where false positives due to colocalization are a concern.

## Acknowledgments

PK and BP were supported by U.S. National Institute of Health grants R01 HG009120. PK was in addition funded by P01 CA134294. RB was supported by the U.S. National Cancer Institute grant R35 CA197449. AG was supported by U.S. National Institute of Health grants R01 GM105857. The breast cancer genome-wide association analyses were supported by the Government of Canada through Genome Canada and the Canadian Institutes of Health Research, the ‘Ministère de l’Économie, de la Science et de l’Innovation du Québec’ through Genome Québec and grant PSR-SIIRI-701, The National Institutes of Health (U19 CA148065, X01HG007492), Cancer Research UK (C1287/A10118, C1287/A16563, C1287/A10710) and The European Union (HEALTH-F2-2009-223175 and H2020 633784 and 634935). All studies and funders are listed in Michailidou et al (2017). The computations in this paper were run on the Odyssey cluster supported by the FAS Division of Science, Research Computing Group at Harvard University.

